# RNA Structure Refinement using the ERRASER-Phenix pipeline

**DOI:** 10.1101/001461

**Authors:** Fang-Chieh Chou, Nathaniel Echols, Thomas C. Terwilliger, Rhiju Das

**Keywords:** RNA structure, structure prediction, X-ray crystallography, refinement, force field

## Abstract

The final step of RNA crystallography involves the fitting of coordinates into electron density maps. The large number of backbone atoms in RNA presents a difficult and tedious challenge, particularly when experimental density is poor. The ERRASER-Phenix pipeline can improve an initial set of RNA coordinates automatically based on a physically realistic model of atomic-level RNA interactions. The pipeline couples diffraction-based refinement in Phenix with the Rosetta-based real-space refinement protocol ERRASER (Enumerative Real-Space Refinement ASsisted by Electron density under Rosetta). The combination of ERRASER and Phenix can improve the geometrical quality of RNA crystallographic models while maintaining or improving the fit to the diffraction data (as measured by R_free_). Here we present a complete tutorial for running ERRASER-Phenix through the Phenix GUI, from the command-line, and via an application in the Rosetta On-line Server that Includes Everyone (ROSIE).

## 1. Introduction

Over the last decade, fruitful progress in RNA X-ray crystallography has revealed three-dimensional all-atom models of numerous riboswitches, ribozymes, and ribonucleoprotein machines (1–3). Due to the difficulty of manually fitting RNA backbones into experimental density maps, many of these crystallographic models contain myriad unlikely conformations and unusually close contacts as revealed by automated MolProbity tools for geometric evaluation of models (4, 5). Inspired by recent advances in *ab initio* RNA structure prediction (6–8) and successful applications of the Rosetta modeling suite in crystallographic and electron microscopy density fitting problems (9, 10), we recently developed the ERRASER method and coupled it with Phenix diffraction-based refinement (11) into a pipeline. In our previous publication (12), we demonstrated that the ERRASER-Phenix pipeline resolves the majority of steric clashes and anomalous backbone and bond geometries assessed by MolProbity in a benchmark of 24 RNA crystal structures. Furthermore, this method led to models with similar or better R_free_. This chapter describes the details of using ERRASER in three easily accessible ways: by a GUI in the Phenix package, from the command-line, and with the ROSIE server (13).

## 2. Materials

The ERRASER-Phenix pipeline relies on two software toolkits: the Rosetta modeling suite (14) and the Phenix package (11). These two toolkits are currently officially supported on Linux and Mac-OSX platforms. (Phenix is available on Windows; Rosetta might be compiled in Windows using Cygwin but is not official supported and well-tested.) To run the pipeline locally, the user needs to have the following versions of the above toolkits installed on their computer:

Rosetta (version 3.5 or later) http://www.rosettacommons.org/
Phenix (version 1.8.3 or later) http://www.phenix-online.org/

Both Rosetta and Phenix are freely available to academic and non-profit institutions. Details of downloading, licensing, and the installation instructions can be found in the above listed websites. Phenix installation procedures can be found at http://www.phenix-online.org/documentation/install.htm. On a Mac-OSX platform installation simply consists of downloading a .dmg file and double-clicking the icon. On Linux systems it consists of unpacking a tar archive and running an installation script. Procedures for Rosetta installation compatible with Phenix and ERRASER can be found at http://www.phenix-online.org/documentation/erraser.htm.

It also possible to run the ERRASER part of the pipeline online and privately using the ROSIE server (http://rosie.rosettacommons.org/).

## 3. Methods

The standard ERRASER-Phenix pipeline consists of three major stages: an initial Phenix refinement, followed by iterative ERRASER refinement, and a final Phenix refinement (Fig. 1). Here the initial Phenix refinement can be skipped if the input structure has already been refined with all hydrogen atoms included in the model. In general, we find that maintaining hydrogen atoms during diffraction-based refinement tends to give models with better geometrical quality, particularly with regards to steric interactions as assessed by the MolProbity clashscore. Since ERRASER performs only real-space refinement, a final diffraction-based refinement is necessary to fit the model directly to the original data and evaluate the R_free_ statistics. We have carried out extensive tests using the Phenix refinement tool for these two refinement stages (15), but users should be able to substitute in refinement tools if preferred (e.g. SHELXL (16), Refmac (17), CNS (18), etc.).

**Figure 1.**
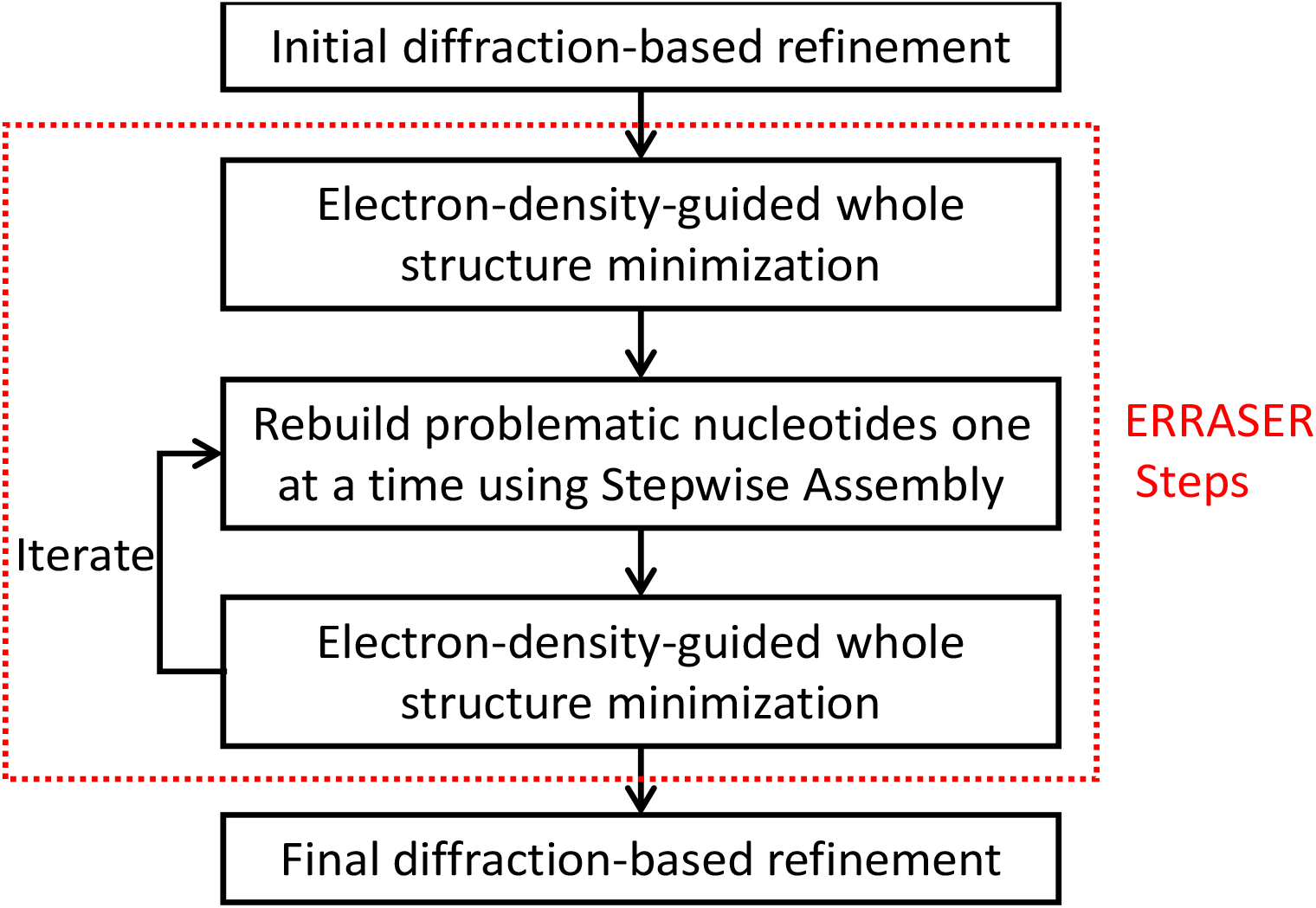
Flow chart of the ERRASER-Phenix pipeline.

In the sections below, we will focus on the details of the ERRASER refinement stage. We will mainly discuss how to run ERRASER using the Phenix GUI interface, and discuss how to run ERRASER using shell command lines and ROSIE web server. Finally, we briefly discuss some settings and options we found useful in the Phenix refinement of RNA.

### 3.1. Set up the Phenix-Rosetta connection

After both Phenix and Rosetta are properly installed and compiled on the user’s local computer, the user should set the path so that Phenix can locate the Rosetta applications. Suppose you have Rosetta installed at “/home/user/rosetta-3.5”. If using the bash or sh shells, add the following line into “∼/.profile” or “∼/.bashrc”:

~~~
export PHENIX_ROSETTA_PATH=/home/user/rosetta-3.5
~~~

Or if using C-shell, put the following line into “∼/.cshrc”:

~~~
setenv PHENIX_ROSETTA_PATH /home/user/rosetta-3.5
~~~

### 3.2. Prepare the ERRASER input files

The following files need to be prepared before running ERRASER. Note that ERRASER is designed for the final fine-tuning of the RNA models, and has only been tested for such cases. Applying ERRASER to models from earlier refinement stages might lead to unexpected behavior and poor models.

#### 3.2.1. Input PDB file

ERRASER accepts PDB-formatted files of RNA with the standard format as input. The users should ensure the input file follows the standard PDB format, with correct residue and atom naming conventions. In addition, the user should ensure that there are no missing heavy atoms in the model, or ERRASER may not run properly.

#### 3.2.2. Density map file

Currently ERRASER supports input 2*mF*_o_-*DF*_c_ density maps in the CCP4 format. It is important to exclude R_free_ diffraction data during map creation to ensure that ERRASER is not influenced by the set-aside data and that final R_free_ values are appropriate for cross-validation. All density data covering the entire unit cell should be included to allow ERRASER correctly evaluate the correlation to density during the sampling. Here we demonstrate the process of creating the density map using the “calculate maps” GUI in Phenix (19). In the GUI window, click the “CCP4 or XPLOR Maps” button, and a window will pop out. In the new window, select the following options (Fig. 2):

~~~
Map type: 2mFo-DFc
File format: CCP4
Map region: Unit cell
Enable the following three options:
        1. “Kicked”
        2. “Fill missing f obs”
        3. “exclude free r reflections”
~~~

**Figure 2.**
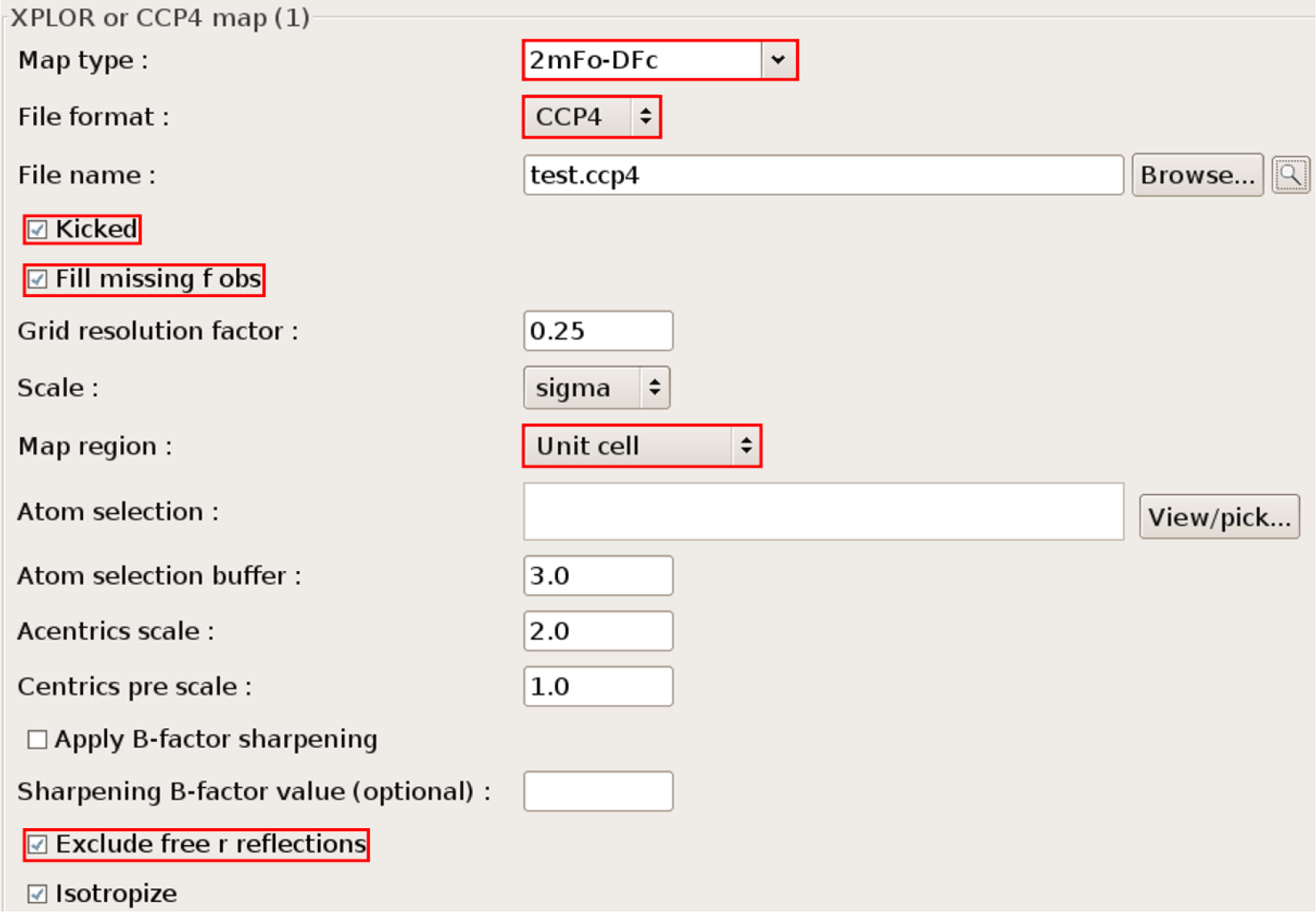
Snapshot and useful options of the phenix.maps utility.

Other options are kept default. Here the “kicked” option makes use of the kicked map algorithm to improve the map quality and to reduce map bias (19). The “Fill missing f obs” option will allow Phenix to fill missing experimental diffraction data (*F_obs_*) with calculated diffraction (*F_calc_*), thereby avoiding the Fourier truncation error. We found that these two options lead to better ERRASER results (e.g., in terms of final *R*_free_) in many cases. “Exclude free r reflections” ensures that the test set of reflections is not included. If you fail to check this option then your final *R*_free_ may be biased and be misleadingly low.

### 3.3. A simple ERRASER job example

It is straightforward to run ERRASER using the Phenix GUI. Click on “Refinement”-> “ERRASER” in the main Phenix GUI to access the ERRASER GUI (Fig. 3). As a quick start, put in a PDB file and a corresponding CCP4 map using the “Input PDB” and “CCP4 map” boxes. Then click on the “Run” button. ERRASER will run in a new tab. Descriptions of ERRASER options that the user may wish to explore are given below in section 3.5.

**Figure 3.**
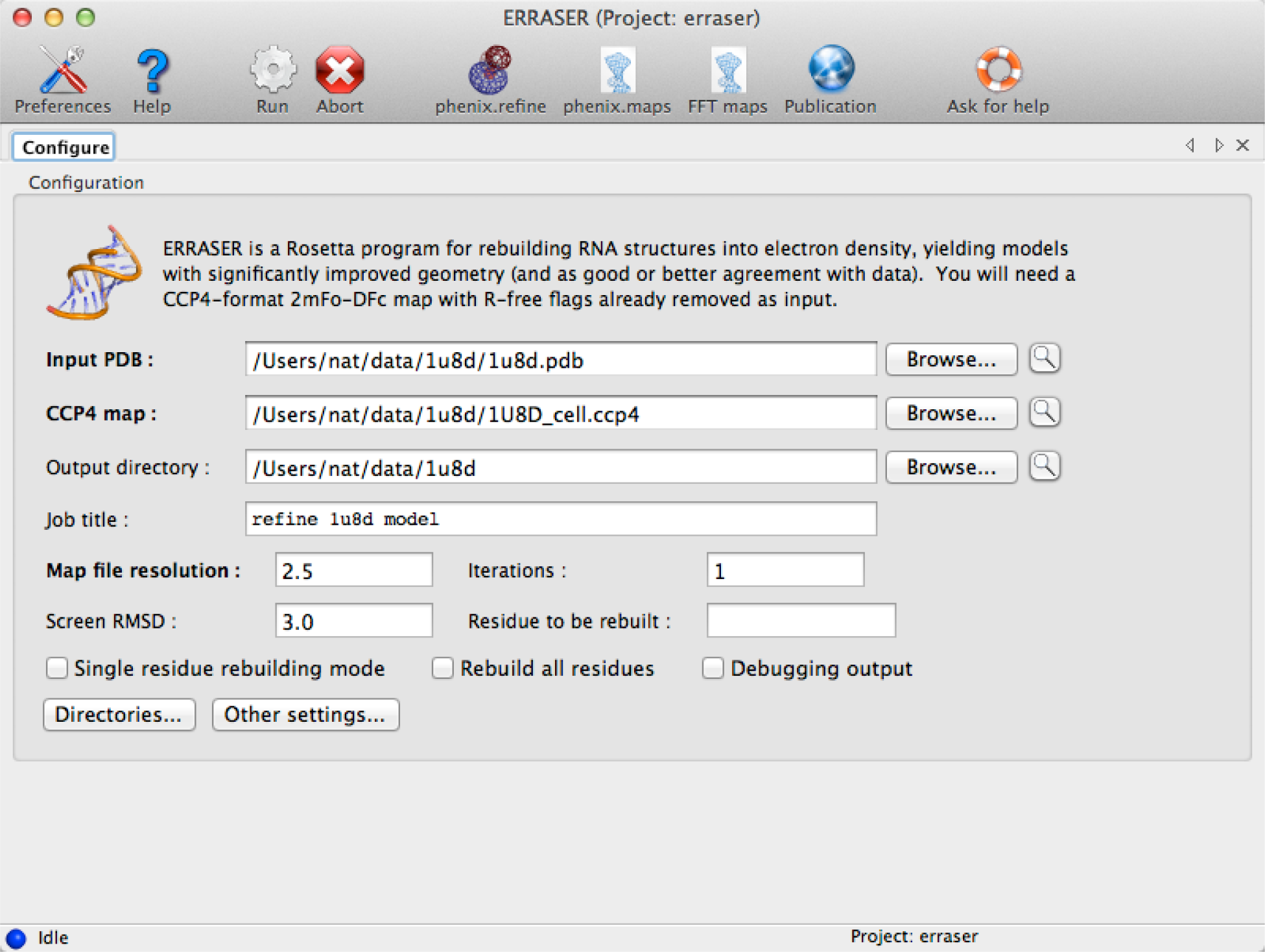
The main window of the Phenix ERRASER GUI.

**Figure 4.**
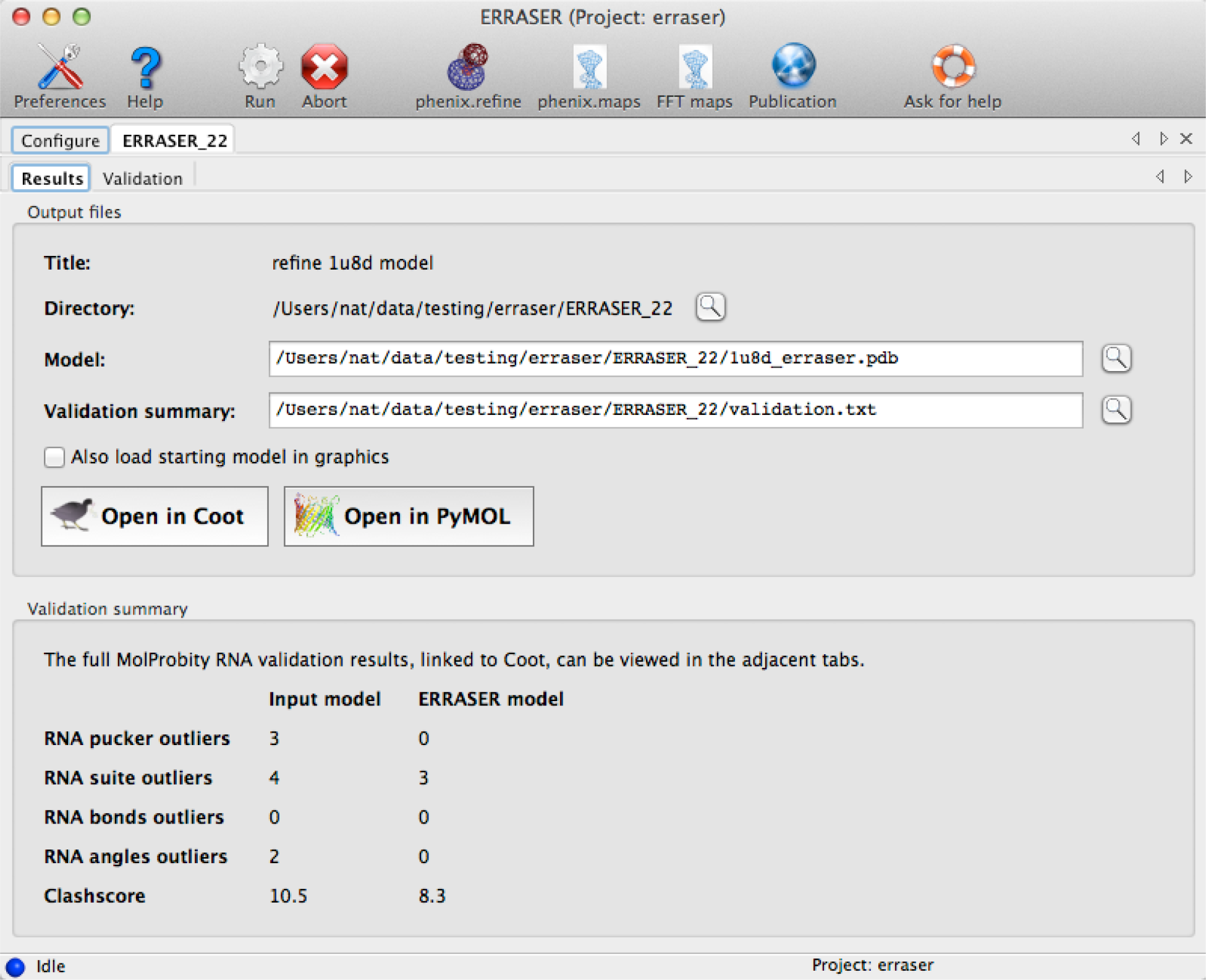
Output window of the ERRASER GUI.

After refinement is complete, ERRASER runs several validation metrics from MolProbity (4, 5) as implemented in Phenix. Results are summarized along with output files in a new tab (Figure 5), with buttons to load the results in Coot and PyMOL. The validation display is identical to the all-atom contact and RNA-specific components in the Phenix GUI, which interact directly with Coot (20). These include:

– Steric clashes (21), defined as atomic overlaps of at least 0.4 Å when explicit hydrogen atoms are present;
– RNA bond length and angle geometry outliers, which has values > 4 s.d. from the Phenix reference values.
– RNA pucker outliers. RNA sugar rings typically adopt either C2′-endo or C3′-endo conformation, known as the pucker. MolProbity can confidently identify erroneous puckers by measuring the perpendicular distance between the 3′-phosphate and the glycosidic bond vector.
– RNA ‘suite’ outliers. Previous research (22, 23) shows that most of the RNA backbone ‘suites’ (sets of two consecutive sugar puckers with 5 connecting backbone torsions falls into 54 rotameric classed. Non-rotameric outlier ‘suites’ are likely to be problematic.

**Figure 5.**
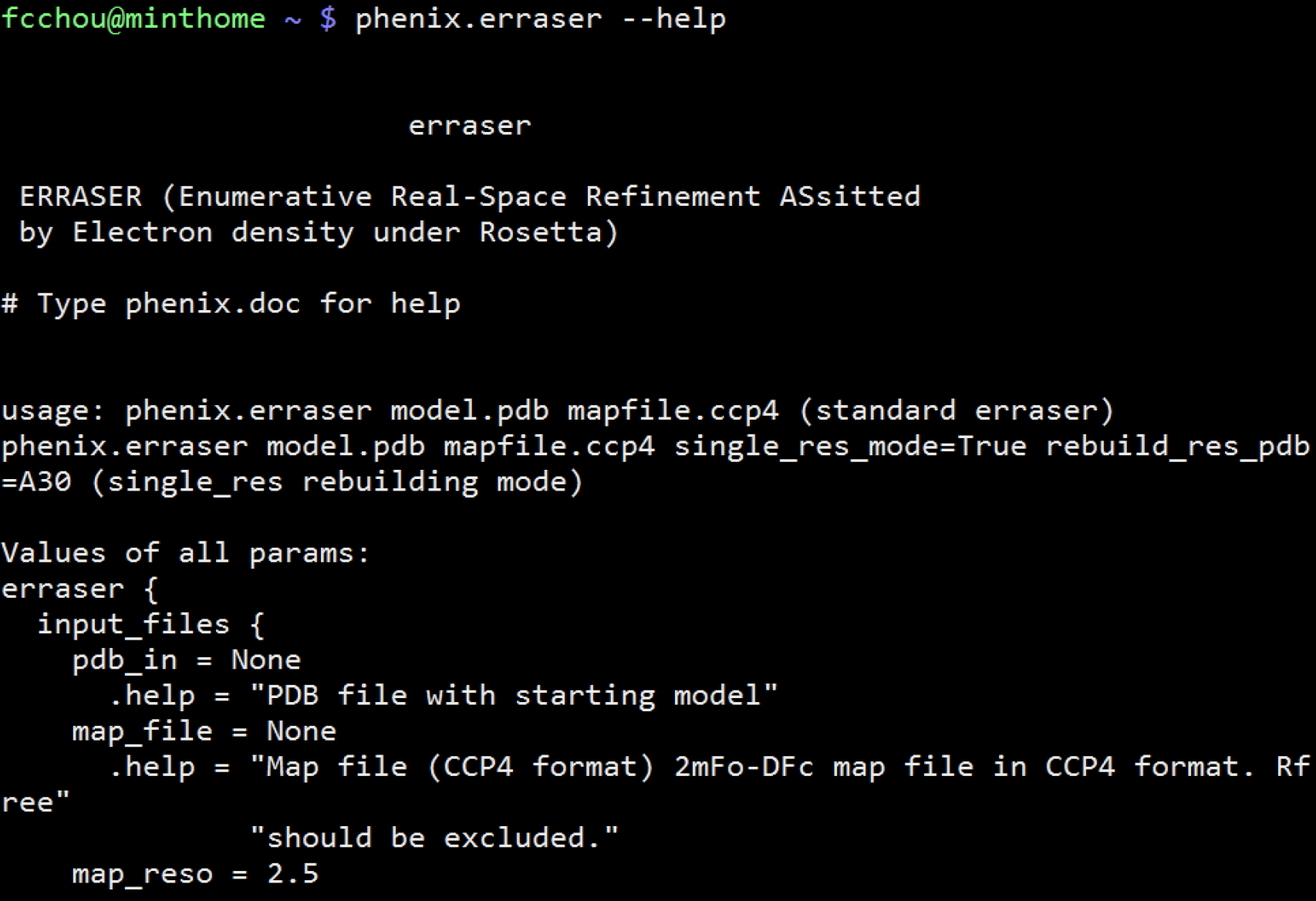
The phenix.erraser command line application.

Phenix also outputs a “Validation summary” text file that compares the model before and after the ERRASER refinement, which can be opened from the GUI by clicking on the corresponding button. In addition to the standard MolProbity metrics above, the “Validation summary” also gives a list of χ angle (glycosidic bond torsion) flips by ERRASER (see Note 1).

The users can use this “Validation summary” list to guide their manual inspection of the post-ERRASER model. If during the inspection, the users think that any residue is not properly rebuilt in the standard ERRASER run, they might try to rebuild such residue with the single-residue rebuilding mode described below.

### 3.4. Single-residue rebuilding mode

For model regions with low density or for which other information is known (e.g., based on homology to a high-resolution structure), ERRASER’s top picks for residue conformations may not be optimal. In addition, ERRASER might not always resolve all errors in the models and might, in a few cases, lead to new errors in the model while fixing other errors. In such cases, the user may want to inspect other possibilities for residue conformations. Therefore, in addition to the standard ERRASER protocol described above, there is a single-residue rebuilding mode available in ERRASER. In this mode, the user selects a particular residue in the complete structure and rebuilds it using ERRASER algorithm. Up to ten different models, sorted by their relative ERRASER score, are returned as output which the user can than inspect.

The users can run the single-residue mode under the same GUI interface similar to the standard ERRASER application. After inputting the PDB file and CCP4 map file, click on the “single residue rebuilding mode” checkbox, and input the residue to be rebuilt in the “Residue to be rebuilt” box. The format of input residue is chain ID followed by residue number, e.g. ‘A35’ or ‘F152’. Then click “Run” button to execute the program. After completion, the generated models and the validation results are displayed in the summary tab. The validation includes the ‘suite’, pucker, and glycosidic torsion assignments, as well as the final ERRASER scores.

### 3.5. Available options in ERRASER

The following options can be found in the main window of ERRASER GUI:

#### 3.5.1. Map file resolution

It is usually a good idea to provide the resolution of the input density map, so that ERRASER can give a more accurate estimation of the fit between the model and the input map. If not provided, ERRASER will assume a default value of 2.5 Å.

#### 3.5.2. Iterations

The total number of iterations of the ERRASER cycles (Fig. 1). By default it is 1. The user can increase the number to perform multi-cycle ERRASER refinement. We recommend not to use any value greater than 3. Beyond 3 iterations, the ERRASER model will most likely be converged, and it is a waste of time to continue the iterations.

#### 3.5.3. Screen RMSD

To speed up the ERRASER rebuilding process, by default ERRASER only considers conformations within 3.0 Å RMSD of the input model during the sampling of each residue. The user can change this cutoff RMSD value by editing this option. If the screen RMSD is set to be larger than 10.0 Å, ERRASER will simply skip the RMSD screening step.

#### 3.5.4. Residue to be rebuilt

This is for the single-residue rebuilding mode only. See section 3.4.

#### 3.5.5. Single residue rebuilding mode

This is for the single-residue rebuilding mode only. See section 3.4.

#### 3.5.6. Rebuild all residues

By default ERRASER will only rebuild residues that are assessed by Molprobity as having outlier pucker, suite, bond length or bond angle, or residues that moved significantly during the Rosetta minimization step. Selecting this option enforces ERRASER to rebuild all residues in the model. In general, however, we recommend using the default rebuilding protocol. Rebuilding all residues is likely to be a time-consuming process without obvious model improvement. In some cases such excessive rebuilding might even lead to worse models, due to artifacts in the input density map and the Rosetta scoring function. If the user wishes to rebuild some specific residues, use the “Extra residues to be rebuilt” option below.

#### 3.5.7. Debugging output

This option is for debugging only. Turning on this option will generate a lot of output and slow down the code. If you find a bug, it might be a good idea to run this mode and send all the output to the developers to aid diagnosis.

The following options are accessible by clicking the “Other settings” button in the GUI.

#### 3.5.8. Fixed residues

This option allows user to specify residues that should be kept fixed throughout the ERRASER refinement. The format is “A32”, where A is the ID of the chain and 32 is the residue number. The user can also input “A22-30” to fix all residues from A22 to A30. This option is especially useful when the RNA contains ligands, modified nucleobases, and strong crystal contacts, all of which are not currently modeled in ERRASER. In these cases, residues near to these unmodeled part of the molecules might get rebuilt into unreasonable conformations due to missing of critical interactions to the unmodeled parts. It is therefore necessary to fix these residues using the “fixed residues” option. See Note 2 for general suggestions on determining the necessary fixed residues.

#### 3.5.9. Extra residues to rebuild

This option allows user to specify particular residues in the model that ERRASER must rebuild regardless whether it has apparent errors. The format is the same as “fixed residues” above.

#### 3.5.10. Native syn only pyrimidine

Syn conformers of pyrimidines (U and C) are rare in RNA structures. By default, ERRASER only samples syn conformers of pyrimidine residue if the user supplies an input residue with the syn conformer. The purpose is, first, to speed up the computation, and second, to avoid possibly problematic syn pyrimidine conformers that show up in the final models due to artifacts of the electron density map. Turning off this option will allow ERRASER to sample syn pyrimidine in all cases. In practice we found that this option is advantageous for low-resolution models (> 2.5 Å, as a rule of thumb); for high-resolution models where it is possible to distinguish syn/anti conformers from the density map, the user may explore turning off this option. The users should carefully examine all the syn-anti pyrimidines flips in the final model if this option is turned off, to make sure there are no suspicious syn pyrimidines in the models (Note 1).

#### 3.5.11. Constrain chi

By default, ERRASER applies a weak constraint on the χ angle (glycosidic torsion) to favor χ conformers that are similar to the input model. The purpose is also to avoid suspicious syn-anti flips during rebuilding; only alternative χ conformers with obvious energy bonuses will be accepted into the final model. Similar to “native syn only pyrimidine” option above, this option is worth exploring for high-resolution data sets, but the user should carefully examine the resulting output (Note 1).

The following options are accessible by clicking the “Directories” button in the main GUI.

#### 3.5.12. Rosetta path

If you have not followed section 3.1 to set up the PHENIX_ROSETTA_PATH environmental variable, you can enter the path to root Rosetta folder here. Otherwise just leave it blank.

#### 3.5.13. Rosetta eraser directory name

This option specifies the location of ERRASER scripts in Rosetta root folder. By default Phenix should automatically figure out the correct path for ERRASER, but if the folder organization in future Rosetta release changes, the user can modify it here to make Phenix and ERRASER compatible.

### 3.6. Run ERRASER from command line

It is also easy to run ERRASER from command using the phenix.erraser command line tool (Fig. 5). As a simple example, just run

~~~
phenix.erraser model.pdb map.ccp4
~~~

Here “model.pdb” is the input pdb model, and “map.ccp4” is the density map file. For a complete list of all available options in phenix.erraser, simply run

~~~
phenix.erraser --help
~~~

For example, one can use the following command line:

~~~
phenix.erraser model.pdb map.ccp4 single_res_mode = True rebuild_res_pdb = A25 map_reso = 2.0
~~~

This allows you to run ERRASER in the single-residue mode to rebuild residue A25. The input map resolution is also given as 2.0 Å.

### 3.7. Runing ERRASER using the ROSIE server

An alternative way of running ERRASER is to use to ROSIE online server (Fig. 6). In this way you do not have to install Phenix and Rosetta locally in your computer. The ROSIE ERRASER app can be found at http://rosie.rosettacommons.org/erraser/. Simply follow the online documentation to run the job. ROSIE can run ERRASER in both standard mode and single-residue rebuilding mode. Note that the ROSIE server has fewer tunable options than local installation. For example, users cannot currently carry out multiple iterations of ERRASER run (it is still possible manually; see Note 3). One additional warning: the memory on each core of ROSIE backend is limited; therefore the job may crash if the user inputs a very large electron density map file in ROSIE.

**Figure 6.**
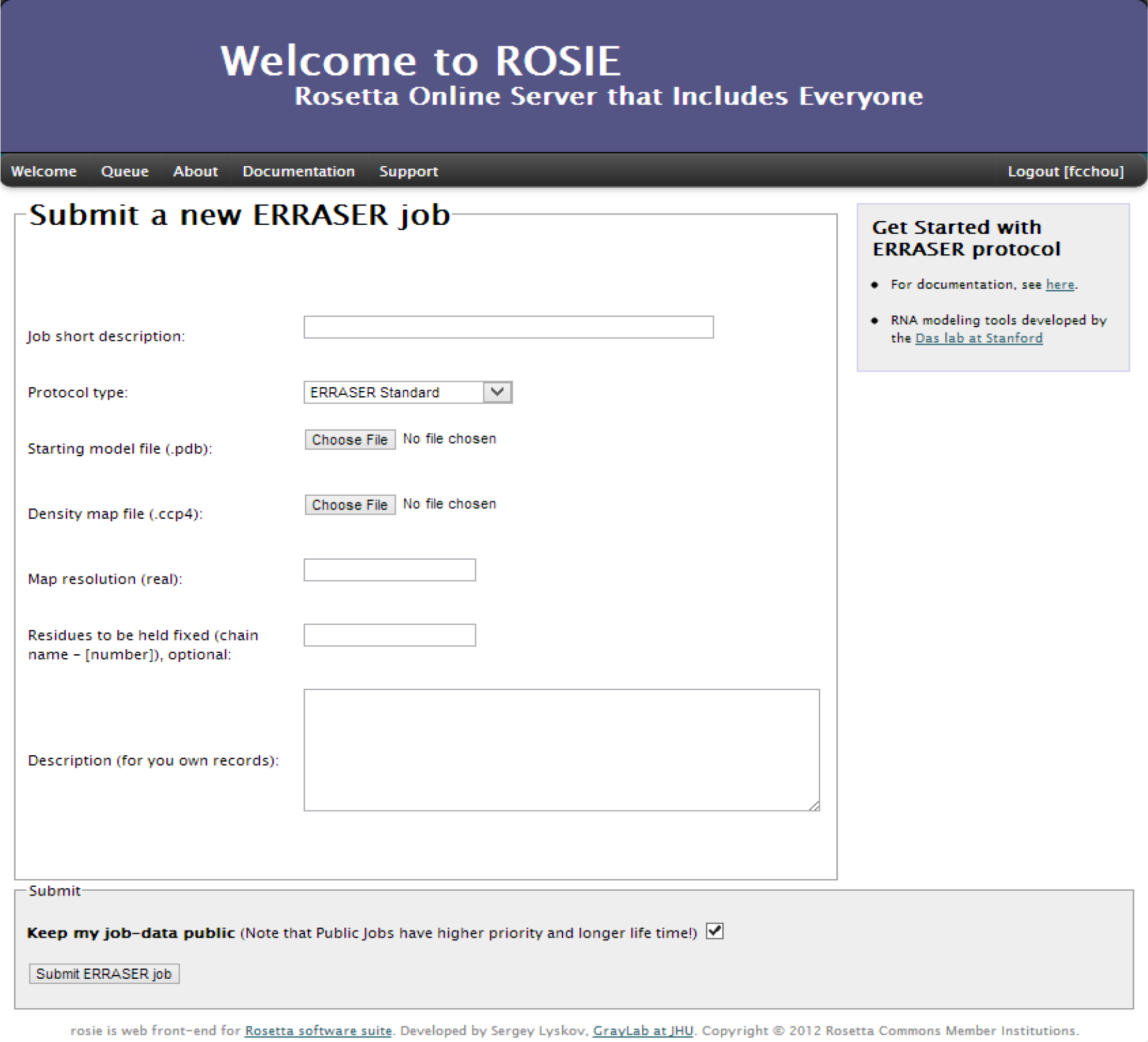
The ROSIE ERRASER cloud application.

### 3.8. Advice for Phenix RNA refinement

The refinement tool in Phenix can be accessed by the phenix.refine GUI from the main Phenix window. A comprehensive documentation for the phenix.refine GUI can be found in the Phenix website (http://www.phenix-online.org/documentation/refinement.htm). Here we will discuss some non-default options that we found useful in refining RNA crystallographic models produced by ERRASER. All of the options discussed below can be found under the “refinement setting” tab in the phenix.refine GUI. Since the refinement setting is sensitive to the initial model and diffraction data, there is no generic rule in setting up the refinement that works in all cases. The user can monitor the change of R, R_free_ and other geometric validation results of the refinement outputs to help decide whether the selected refinement strategy is appropriate.

#### 3.8.1. Refinement strategy: Real-space

We recommend turning off this option during refinement of ERRASER models. By default Phenix alternates between real-space and diffraction-based refinement during a refinement cycle. As the ERRASER models have already gone through extensive real-space refinement in the ERRASER protocol, further real-space refinement in Phenix usually leads to no model improvement, or even leads to worse models in terms of clashes or other quality measures. For the exact parameters being used in refining the models in the 24 ERRASER benchmark set, please refer to the supplementary information of the original ERRASER paper.

#### 3.8.2. Refinement strategy: Number of cycles

While the default of three cycles of refinement is usually good enough to refine the ERRASER models, sometimes more cycles of refinement leads to better results. The user may try a few different numbers (usually 3 to 10) for best results.

#### 3.8.3. Targets and weighting: Optimize X-ray/ADP weights

This option enables Phenix to perform a grid search to find the best X-ray/ADP (Atomic Displacement Parameter, also known as B-factor) weight ratio during each refinement cycle. In practice we found using this option generally leads to final models with better R and R_free_.

#### 3.8.4. Targets and weighting: Optimize X-ray/Stereochemistry weights

This option enables Phenix to perform a grid search to find the best X-ray/Stereochemistry weight ratio during each refinement cycle. In some cases this option leads to better R and R_free_, but in other cases it is better to use a constant X-ray/ Stereochemistry weight (see below). For best refinement results, the user can try both and select the best model.

#### 3.8.5. Targets and weighting: Refinement target weights: wxc_scale

This option sets constant X-ray/Stereochemistry weight during the refinement. In practice we found that it is better to use a lower wxc_scale for models with lower resolution. For example, we used the following heuristic rule in determining the wxc_scale in refining the ERRASER benchmark: wxc_scale = 0.5 for Resolution < 2.3 Å, wxc_scale = 0.1 for 2.3 Å ≤ Resolution < 3 Å, wxc_scale = 0.05 for 3 Å ≤ Resolution ≤ 3.6 Å, and wxc_scale = 0.03 for Resolution > 3.6 Å.

#### 3.8.6. Other options: update waters

The option enables automatic water picking during Phenix refinement. This procedure removes crystallographic waters lacking evident electron density and adds waters to locations with strong unoccupied electron density. In general we found this algorithm improved the model and reduced R and R_free_.

## 4. Notes

1. ERRASER can introduce flips of the χ angles (glycosidic torsion) during the run; we have shown previously that many of these flips are accurate when comparisons can be made of ERRASER-refined models to higher resolution data sets. If “constrain chi” or “native syn only pyrimidine” options are turned off, χ angle flips are made more often, and some of these flips may be incorrect. In low-resolution density map (> 2.5 Å), it is hard to determine the glycosidic conformer by the density map, especially for pyrimidines. While most χ angle flips are reasonable, there are cases where the flips are likely to be incorrect. We recommend that the user to inspect visually all the χ angle flips before proceeding with the final model. Here are some general suggestions on examining the χ angle flips: First, syn conformers are rare, especially for syn pyrimidines. Any anti-to-syn flips should be closely examined. Unless there is strong electron density evidence or new hydrogen bond interactions, the flips are likely to be problematic. Syn-to-anti flips are more likely to be correct, but the user should still examine the models to make sure no important hydrogen-bonding interactions are broken during the flips.
2. Currently ERRASER does not handle crystal contact, ligands and modified residues during the refinement. To address these issues, the user can use the “fixed residues” option. Residues contacting ligands and residues in crystal contacts can be fixed by adding them to the list of “fixed residues”. For modified residues, one should further constrain the residues which the modified residue directly bonded to. For circular RNA, one can constrain both the first and the last residue to ensure the bond does not break during ERRASER refinement. An alternative way to handle crystal contacts is to manually add in the crystal packing partners during ERRASER step; this can be achieved using the symexp utility in PyMol. Usually it is unnecessary to add in the entire molecules of the crystal packing partners; one can cut out just the relevant residues nearby. These extra add-in residues should be removed from the final ERRASER models before Phenix refinement.
3. To manually iterate ERRASER refinement in ROSIE, simply use the output pdb file of the first job as the input pdb file for a second job, and use the same input density map and options as the first iteration.

## Acknowledgements

The authors would like to thank Sergey Lyskov for implementing and maintaining the ROSIE Rosetta server, and members of the Rosetta and the Phenix communities for discussions and code sharing. The authors acknowledge support from Howard Hughes International Student Research Fellowship (F.C.), Stanford BioX graduate student fellowship (F.C.), Burroughs-Wellcome Career Award at Scientific Interface (R.D.), NIH grants R21 GM102716 (R.D.) and P01 GM063210 (P.D. Adams, PI, to N.E. and T.T.).

